# Network effects of traumatic brain injury: from infra slow to high frequency oscillations

**DOI:** 10.1101/2023.08.31.555809

**Authors:** Brianna Marsh, Mingxiong Huang, Maxim Bazhenov

## Abstract

Traumatic brain injury (TBI) can have a multitude of effects on neural functioning. In extreme cases, TBI can lead to seizures both immediately following the injury as well as persistent epilepsy over years to a lifetime. However, mechanisms of neural dysfunctioning after TBI remain poorly understood. To address these questions, we analyzed experimental data and developed a biophysical network model implementing effects of ion concentration dynamics and homeostatic synaptic plasticity to test effects of TBI on the brain network dynamics. We focus on three primary phenomena that have been reported *in vivo* after TBI: an increase in infra slow oscillations (<0.1 Hz), increase in delta (0.1 - 4 Hz) power, and the emergence of high frequency oscillations (HFOs) in the gamma range (30 - 100 Hz). We show that the infra slow oscillations can be directly attributed to extracellular potassium fluctuations, while the existence and characterization of HFOs is related to the increase in strength of synaptic weights from homeostatic synaptic scaling. The experimentally found transient increase in delta power can be attributed to the inter-HFO timings. We then show that buildup of high frequency oscillations in the injured region can lead to seizure-like events that span all neurons in the network; additional seizures can then be initiated in previously healthy regions. This study brings greater understanding of network effects of TBI, and how they can give rise to epileptic activity. This lays the foundation to begin investigating how injured networks can be healed and seizures prevented.

**Significance Statement:** This project delineates and attempts to explain abnormalities seen in human brain following traumatic brain injury (TBI). TBI can lead to the development of seizures, which may last a lifetime and often become resistant to pharmaceutical treatments. The study identified key mechanisms responsible for occurrence of three characteristic changes in spatio-temporal network dynamics following TBI. This model provides predictions that can serve as a testing ground for potential therapeutic approaches.

## Introduction

Traumatic Brain Injury (TBI), while often spoken of as a monolith, can come in many different forms and from many different sources, such as rotational shearing, blunt force or incision, or internal bleeding(Saatman et al., 2008). What they all have in common, however, is the ability to disrupt normal neural functioning and activity. Some TBI’s can cause abnormal synchrony that escalates to seizures both immediately following the trauma and after a long time delay – months or even years years later(Avramescu and Timofeev, 2008; Chauvette et al., 2016; Dinner, 1993). While treatments exist for epilepsy, up to 30% of patients are drug resistant in the long term(Łukawski et al., 2018). Due to this, novel approaches including intervention following injury that addresses abnormal network behavior are needed. Currently, it is a common clinical practice to administer drugs that inhibit neural activity following trauma, in anticipation of possible upcoming seizures(Chartrain et al., 2017; Torbic et al., 2013). However, this is a blunt instrument at best that does not address long term underlying pathology. To address these problems, we first need to fully understand and characterize the injured network behavior. In this paper, we aim to do just that.

Following traumatic brain injury, marked hyperactivity in the Gamma range (30-80 Hz) has been measured in patients with post-concussion syndrome and in task-evoked MEG during working memory tasks(Bailey et al., 2017; Culic et al., 2005; Huang et al., 2020, 2019). Increases in slow oscillations have also been observed in relation to traumatic brain injury and epilepsy(Franke et al., 2016; Guerriero et al., 2022; Huang et al., 2019). Finally, several studies have reported an increase in the Delta range(Davenport et al., 2022; Safar et al., 2021; Sandsmark et al., 2017; Thomasy et al., 2016) activities. These are the three post-injury phenomena we aim to explore and to explain in our study.

We previously proposed that following brain trauma, the sudden lack of input to the injured neurons induces increases in the synaptic weights of remaining connections in an attempt to regain a normal firing rate(Fröhlich et al., 2008; González et al., 2015; Houweling et al., 2005). This regulatory mechanism of synaptic weight changes to maintain network firing rates is called Homeostatic Synaptic Plasticity (HSP)(Burrone and Murthy, 2003; Pozo and Goda, 2010; Turrigiano, 2008). In the extreme case of traumatic brain injury, HSP may fail to correctly compensate for the lost input activity, inducing over-excitation and driving seizures(Fröhlich et al., 2008; González et al., 2019, 2015; Houweling et al., 2005; Timofeev et al., 2013). Here, we hypothesize that HSP-dependent strengthening of connections between cells after TBI may alter network dynamics to generate the highly synchronized bursting activity, which in turn may lead to seizure-like events. We have further previously shown that infra slow oscillations during resting state are strongly tied to potassium fluctuations(Krishnan et al., 2018). Here, we test the hypothesis that HSP-dependent changes in connectivity may explain dramatic increase in these oscillations after injury.

To test these hypotheses and further understand pathological network behavior after injury, we developed a model of traumatic brain injury that includes ion concentration dynamics and HSP. This model displays many phenomena found by analyzing experimental data. We found that following TBI, the network synchronized activity was increased in the infra-slow, delta, and high gamma frequency ranges. This led to spontaneous high frequency bursting events, resembling high-frequency oscillation (HFO) interictal events(Arroyo and Uematsu, 1992; Worrell et al., 2004). When injury was concentrated within a smaller region, these HFO-like events triggered spontaneous spike-and-wave seizures. This study characterized spatio-temporal dynamics of pre-seizure interictal events and revealed underlying mechanisms of their generation in the traumatized brain.

## Materials and Methods

### EXPERIMENTAL DATA ACQUISITION METHODS

Resting-state MEG (rsMEG) data acquisition was measured in alert condition using our Elekta-Neuromag VectorView™ MEG system with 306 MEG channels at the UCSD MEG Center. The entire recording had two 5-minute blocks with eyes closed while staying alert(Huang et al., 2020, 2014b). Data were sampled at 1000 Hz and are run through a high-pass filter [0.1 – 330] Hz cut-off. Eye blinks/movements, and heart signals are monitored.

All study participants were males and U.S. active-duty military service members or Operation Enduring Freedom / Operation Iraqi Freedom Veterans. Twenty-six participants (age 28.2 ± 6.5) had a chronic combat-related blast mTBI and with persistent post-concussive symptoms for an average of 16.2 months post injury (standard deviation = 15.3; range = 4 to 84 months). Combat-related mTBI was corroborated from medical records. Age-matched healthy controls included 19 individuals with similar combat experience (age 2.85 ± 5.9), but without a significant history of concussion based on self-report.

All mTBI participants were evaluated in a clinical interview to assess the nature of their injuries. The diagnosis of mTBI was based in part on standard Veterans Affairs and Department of Defense diagnostic criteria(Management of Concussion/mTBI Working Group, 2009): 1) loss of consciousness < 30 minutes or transient confusion, disorientation, or impaired consciousness immediately after the combat-related trauma; 2) post-traumatic amnesia < 24 hours; and 3) an initial Glasgow Coma Scale(Teasdale and Jennett, 1974) between 13-15 if available. Since the Glasgow Coma assessment was not accessible in combat theater, participants missing an assessment, but who met other inclusion criteria, were also enrolled.

Conventional structural MRI acquisition and the construction of the BEM: Using a 3T GE Discovery MR750 MRI scanner, we will acquire a high-resolution MRI volume with a resolution of 1×1×1 mm3 using a T1-weighted 3D-IRSPGR pulse sequence. Gradient nonlinearity spatial distortion in MRI will be corrected(Jovicich et al., 2006). The T1-weighted images will be used to construct a source grid for Fast-VESTAL and a boundary element method (BEM) model for MEG forward calculation(Mosher et al., 1999). Conventional MRI sequences for identifying TBI lesions in participants will also be performed: 1) Oblique T2*-weighted; 2) Oblique T2-weighted with ASSET; 3) Oblique FLAIR; 4) Oblique DWI. Susceptibility-weighted imaging will also be performed to detect subtle blood products.

### MODEL DESIGN

Our computational model consists of 200 pyramidal neurons and 40 inhibitory neurons. This model is primarily an expansion of the model presented in González (2015)(González et al., 2015); in addition to a larger number of neurons being modeled, HSP is implemented as a local (rather than global) process so that localized effects of injury can be examined. The simulated neurons are conductance-based two compartmental neuron models with realistic ion concentration dynamics, as described in previous publications(Bazhenov et al., 2004, 2002; Fröhlich et al., 2008; González et al., 2015; Houweling et al., 2005; Krishnan et al., 2015a; Krishnan and Bazhenov, 2011; Volman et al., 2011b, 2011a). The membrane potential dynamics for each compartment were modeled by the following equations:

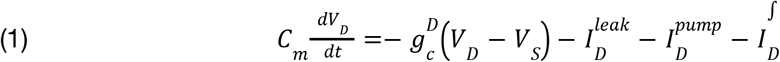

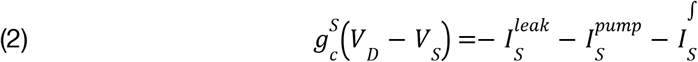

where *V*_*D*_, *V*_*S*_ are the dendritic and axosomatic membrane potentials, 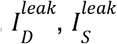 are the sum of the ionic leak currents, 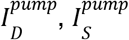 are the sum of the Na+ and K+ currents, and 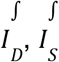 are the intrinsic currents for the dendritic and axosomatic compartments. Reversal potentials for all current are calculated using Nernst equation based on the intra- and extracellular ion concentrations. The intrinsic currents for the dendritic and axosomatic compartments as well as ion concentration dynamics for [K+]o, [K+]i, [Na+]o, [Na+]i, [Ca2+]i, and [Cl−]i have been previously described(González et al., 2015; Krishnan et al., 2015a; Krishnan and Bazhenov, 2011).

The 200 pyramidal and 40 inhibitory neurons are arranged linearly. Each pyramidal cell has a connection radius of 5 to other pyramidal cells (with AMPA conductance strength of 3.5 nS and NMDA conductance of 0.9 nS) and 1 inhibitory cell (with AMPA and NMDA conductance strengths of 2.4 nS and 0.24 nS), while inhibitory cells make a total of 5 unique connections to pyramidal cells (with GABAA conductance strengths of 3.5 nS). This means each pyramidal cell receives input from 10 other pyramidal cells and 1 inhibitory cell. Each neuron additionally receives external Poisson input as baseline activity to simulate long range connections from other cortical regions as described in our previous studies(González et al., 2015; Krishnan et al., 2015a; Krishnan and Bazhenov, 2011).

Homeostatic synaptic plasticity (HSP) is implemented to maintain a firing rate of 5 Hz by adjusting AMPA conductances between excitatory neurons. The relevant firing rate for each neuron is calculated as the mean firing rate of itself and all pyramidal cells within its connection radius. AMPA conductances are then scaled relative to the difference between the current and target firing rates (Equation 2). The HSP scaling factor is here set to 0.01.

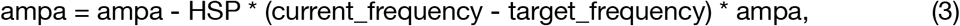

Traumatic Brain Injury is modeled as a reduction to a cell’s Poisson input, to represent lost synaptic connections and broken axons resulting from injury. We here primarily implemented injury as a 50% loss of Poisson input. Injury was applied to a random 40 of the central 80 neurons in the model; these central 80 neurons will be referred to as the Injury Zone. This means that there are 60 healthy neurons on either side of the Injury Zone; these regions will be referred to as the Healthy Zones. We further implemented a more concentrated injury that leads to seizures: in this case, injury was applied to all of the most central 40 neurons, with no healthy neurons interspersed.

This paper identifies and characterizes two types of pathology seen after injury: Infra Slow Oscillations (ISOs) and High Frequency Oscillations (HFOs). ISO’s are here defined as <0.1 Hz oscillations. HFO’s are here defined as sequences of spikes that have an interspike interval less than 33 ms (greater than 30 Hz) for a minimum of 3 consecutive spikes. This was done to exclude single random high frequency spikes. HFO’s often contain much higher frequencies (up to 80 Hz) but this method was chosen to identify as many HFO’s as possible while excluding any behavior seen in the control model or pre-injury timeframes.

Power spectrums of the network are computed using Welch’s method(*PSD computations* using Welch’s method. [Power Spectral Density (PSD)], 1991) with a sliding window of 50 seconds with 50% overlap in order to capture low frequency events. The power spectrum is computed for groups of 5 linearly consecutive neurons, then averaged over groups; this ensures retention of high frequency events, while also getting a spatial average. The local field potential is computed as the smoothed average of all pyramidal neuron voltages.

### EXPERIMENTAL DATA DESIGN & ANALYSIS

Voxel-wise MEG source magnitude images are obtained using our high-resolution Fast-VESTAL method(Huang et al., 2014a). The rsMEG data are first run through MaxFilter(Song et al., 2008; Taulu et al., 2004) to remove external artifacts and correct head movement. Residual artifacts are removed using fast Independent Component Analysis (ICA)(Hyvärinen and Oja, 2000). Artifact-free rsMEG waveforms are divided into 2.5 second epochs with 50% overlaps. Data in each epoch are DC-corrected and then can be run through band-pass filters for different frequency bands, such as delta (1-4 Hz), theta (4-7 Hz), alpha (8-12 Hz), beta (15-30 Hz), and gamma (30-80 Hz). Sensor-waveform covariance matrices are calculated for individual epochs and then averaged across individual epochs. Using the averaged covariance matrix, plus the source grid and BEM model described previously, voxel-wise rsMEG images are obtained for each participant and frequency band using Fast-VESTAL(Huang et al., 2020, 2019, 2014b, 2014a). Voxel-wise MEG images obtained from Fast-VESTAL for rsMEG are spatially co-registered to the Montreal Neurological Institute (MNI-152)(Grabner et al., 2006) brain-atlas template using a standard processing pipeline(Smith et al., 2004). In the present study, we will focus on group analysis of delta-band rsMEG activity.

We analyzed resting state MEG from 26 veterans with blast-related symptomatic mTBI and 19 healthy control veterans with similar deployment experiences. We processed the data as described in the methods to extract the source-magnitude in the delta band, then performed a group analysis of voxel-wise ANOVA that compared rsMEG activity in each band.

In addition, in one representative mTBI and one representative healthy control participants, spectrum analysis based on Fast Fourier Transform (FFT) of the MEG sensor waveform was used to access magnitude differences across frequencies, with the focus on delta (1-4 Hz) and gamma (30-90 Hz) bands. Time-frequency analysis using the Morlet wavelet transformation was also used to illustrate the time-frequency changes of a typical example of abnormal delta- and gamma-band activity from the mTBI participant.

### MODELING DATA ANALYSIS

We first characterized basic features of a control network and an injured network - cell voltage traces, firing rates, and power spectrums. Then, to investigate the effects of potassium and synaptic weights, we applied several different types of manipulations. First, potassium and synaptic weights were frozen, or prevented from further changes at a given point after injury had occurred and pathological behavior had been established. Then, one of the two (potassium or weights) could be set to different levels to investigate the effects of such changes. It is important to note that these experiments require both potassium and the synaptic weights to be frozen before one can be manipulated or set to an extreme; if one is left to vary freely while the other is manipulated, the first will show large compensatory effects. For example, when potassium is set to a high level, the weights decrease correspondingly (supplemental Figure 3). It is then impossible to say if the observed changes are due to high potassium or low synaptic weights. With this procedure in place, we can then determine the role of each in the observed changes in network behavior after injury.

We ran statistical analysis on the observed changes in power across different levels in potassium and synaptic weights. This was done with a simple linear regression using the SciKit-Learn package in Python, with a significance threshold of p < 0.05.

### CODE/SOFTWARE

The computational model was coded in C++; Python was used for subsequent data visualization and statistical analysis.

## Results

### DELTA-BAND VOXEL-WISE MEG SOURCE MAGNITUDE IMAGING: MTBI VS HEALTHY CONTROLS

We analyzed data from 26 veterans with blast-related symptomatic TBI and 19 healthy control veterans with similar deployment experiences. Figure 1 showed the results from the group analysis of voxel-wise ANOVA that compared rsMEG activity in the delta band for 26 veterans with blast-related symptomatic mTBI and 19 healthy control veterans with similar deployment experiences. Abnormal increases were from bilateral hippocampi, frontal poles, dorsolateral prefrontal cortex, superior and middle temporal gyri, primary motor cortex, supplementary motor area, precuneous, and left orbitofrontal cortex.

**Figure 1:**
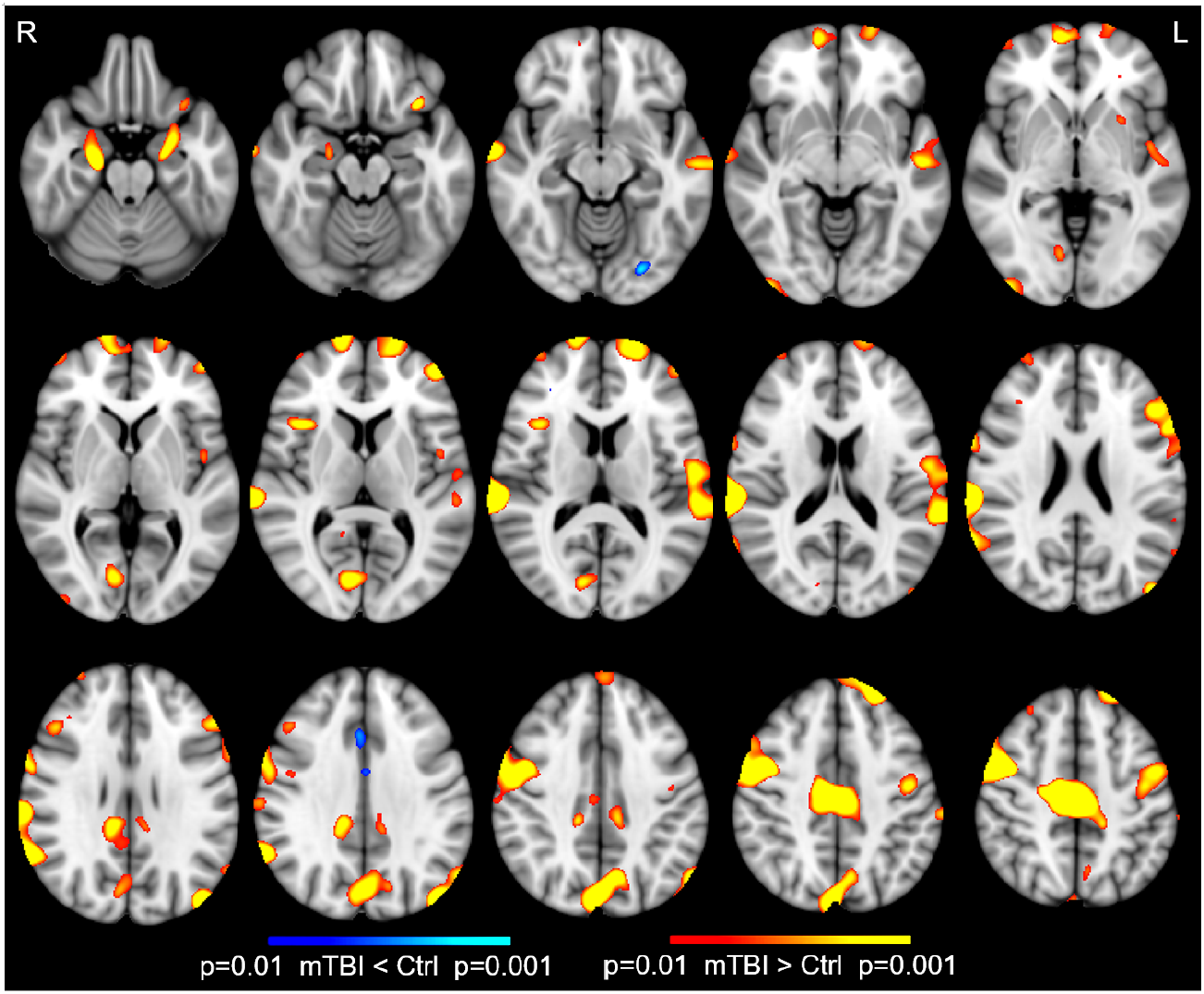
Delta Band Increase Localization. Abnormal increases (regions with red-yellow color) of rsMEG delta-wave (1-4 Hz) activity in 26 blast mTBI veterans, compared with 19 healthy control veterans. Abnormal increases were seen in bilateral hippocampi, frontal poles, dorsolateral prefrontal cortex, superior and middle temporal gyri, primary motor cortex, supplementary motor area, precuneous, and left orbitofrontal cortex.

MEG sensor waveforms from a representative patient with TBI showed an increased slow-wave”burst” (Fig. 2A); the time-frequency plot shows the slow-wave activity was mainly in the delta band and extended to the theta band (Fig. 2B); The MEG spectrum plot during a 5-min recording session from an mTBI and a healthy control subject (Fig. 2C) shows the increase of slow-wave activity mainly in delta (p < 10^−10^) but extending to theta bandwidths. The multiple peaks may be due to more than one generator of abnormal low-frequency activity. These findings showed that blast-related TBI resulted in abnormal increases of delta-band rsMEG activation. The mTBI patient also showed significant increase in the gamma band (excluding the 60 Hz artifact) when compared with the healthy control participant (p < .01), although the difference was not as pronounced as in the delta band.

**Figure 2:**
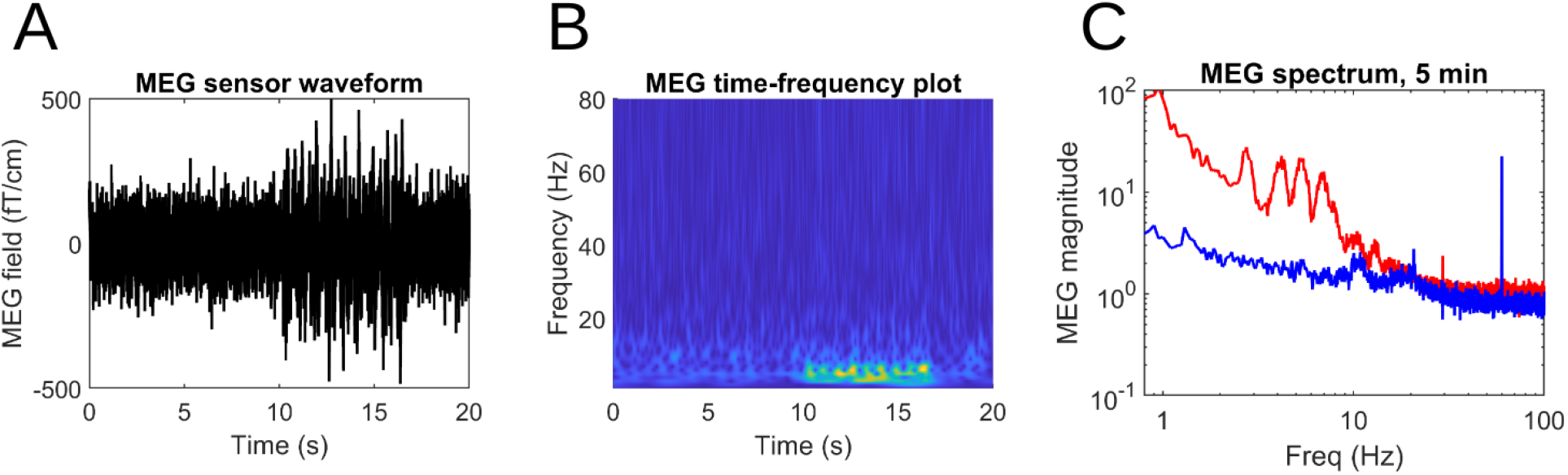
MEG Waveforms. A) rsMEG sensor waveform showing slow waves from one mTBI patient; B) Time-frequency plot of the slow waves; C) Spectra plots of 5-min rsMEG recordings from an mTBI (red) and a control (blue) subject. There are significant increases in both the delta and gamma bands (p < .01). Spike is 60 Hz powerline artifact.

### STEADY RANDOM ACTIVITY IN CONTROL MODEL

The model consists of 200 pyramidal neurons and 40 inhibitory neurons arranged as a linear array; pyramidal neurons have a connection radius of 5, while each inhibitory neuron synapses on to 5 excitatory neurons (Fig. 3). The model neurons are conductance-based two compartment models with realistic ion concentration dynamics (see Methods). Local Homeostatic Synaptic Plasticity (HSP) works to maintain a steady firing rate by adjusting the strength of connections between pyramidal neurons. All neurons further receive steady Poisson input to stimulate long-range connections. Traumatic Brain Injury (TBI) is implemented as a 50% reduction to this long-range input. TBI is applied to a random 40 of the 80 most central neurons to simulate a diffuse injury pattern (Fig. 3). These central 80 neurons will be referred to as the Injury Zone, while the 60 uninjured neurons on either side will be considered the Healthy Zones.

**Figure 3:**
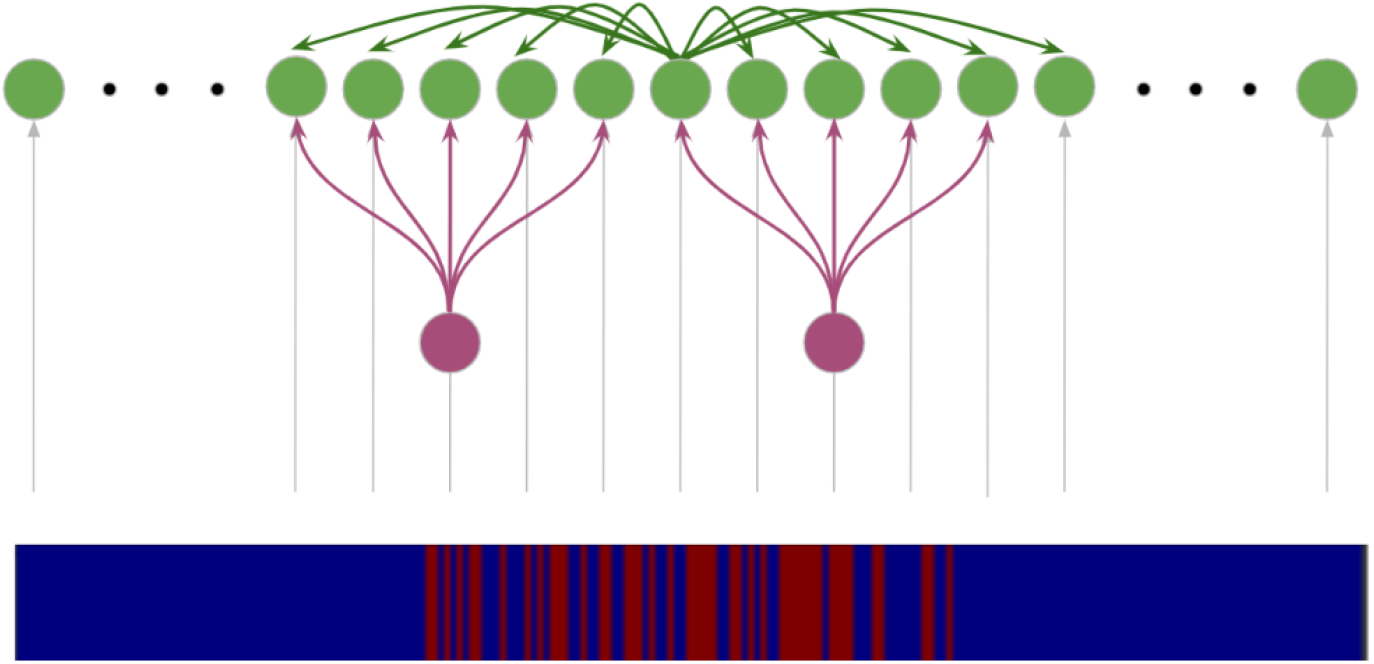
Computational Model Diagrams. (Top) Linear model diagram, depicting each pyramidal cell (green) has a connection radius of 5 to other pyramidal cells (10 total connections). Each inhibitory cell (purple) connects to 5 total pyramidal cells. There are 200 pyramidal and 40 inhibitory neurons in total. (Bottom) Trauma pattern diagram, depicting location of the 40 injured neurons (red lines) across all 200 linear pyramidal neurons.

The model dynamics in a healthy, uninjured state is characterized in Figure 4. The model successfully used local HSP mechanisms to maintain a steady moderate firing rate which is homogenous throughout the network. The power spectral density showed a strong 1/f phenomenon, a hallmark of biological neural data. Single neuron voltage traces showed steady, regular firing while the network local field potential (LFP) revealed very slow low-amplitude oscillations pervasive throughout the simulation.

**Figure 4:**
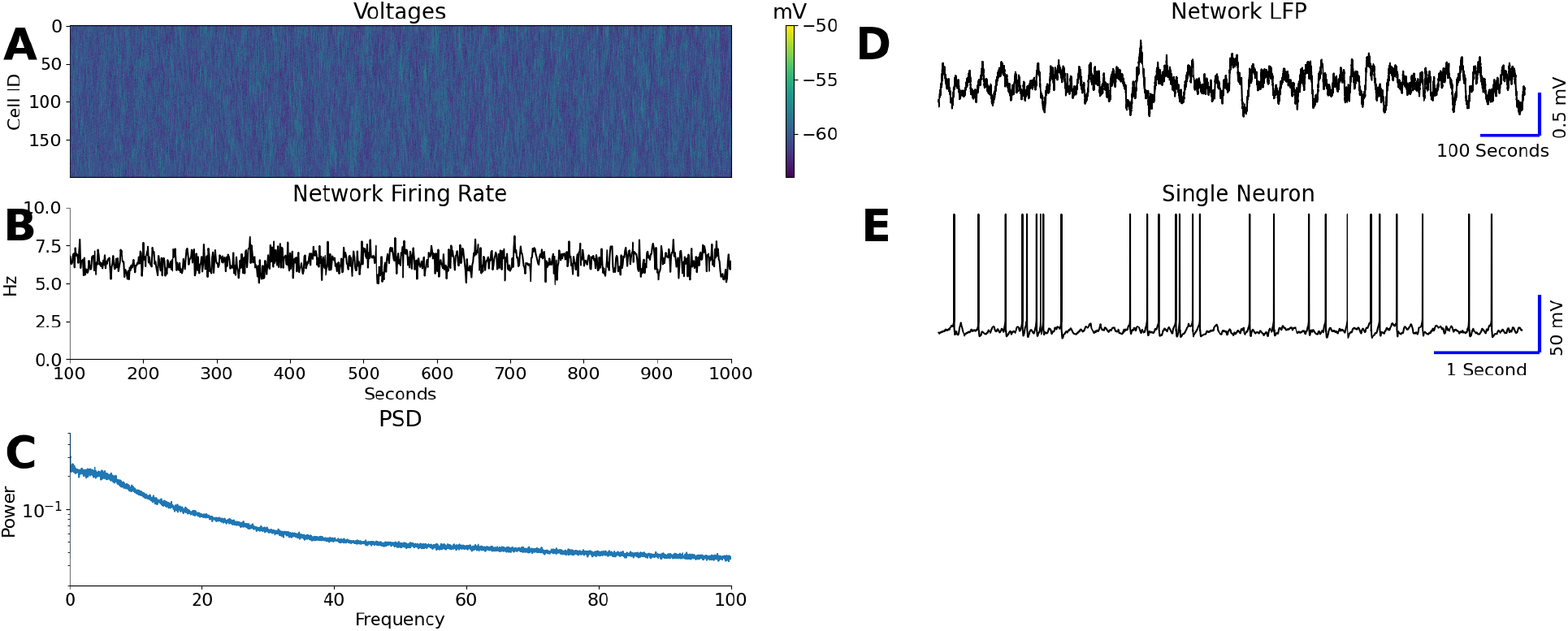
Dynamics of a healthy control network with steady, homogenous activity throughout the network. Raster plot of cell voltages (A) and mean network firing rate (B) are shown for 1000 sec simulation. The power spectral density (PSD) reveals a characteristic 1/f phenomenon (C), while the local field potential (LFP) shows very slow (< 0.1 Hz) oscillations (D). A characteristic single neuron voltage trace in (E) shows spontaneous tonic firing activity.

### SPATIO-TEMPORAL DYNAMICS IN INJURY MODEL

We implemented traumatic brain injury 100 seconds into a simulation as described in the Methods: a random 40 of the central-most 80 neurons had their Poisson input reduced by 50%. These 80 neurons (40 injured neurons interspersed with 40 healthy neurons), here the Injury Zone, are flanked on either side by 60 healthy neurons, here the Healthy Zones. Immediately following this injury, the injured neurons temporarily became silent; the uninjured neurons in the Injury Zone revealed a drop in firing rate, but remained active (Fig. 5A). Because of HSP, the synaptic weights of neurons in the Injury Zone responded to the local drop in firing rate with an increase in the strength of synaptic connections in an attempt to recover the average local firing rate (Fig. 5D). This led to significant increase in amplitude of the infra slow (<0.1Hz) oscillations (Fig. 5B/E) and was mimicked in the potassium concentration (Fig. 5C), which revealed the onset of large fluctuations shortly after injury had occurred. The slow fluctuations were most apparent in the injury zone, but higher potassium did spread out to the Healthy Zones during the fluctuation peak. A voltage trace of a single injured neuron revealed high density bursts of spikes with minimal tonic firing (Fig. 5F).

**Figure 5:**
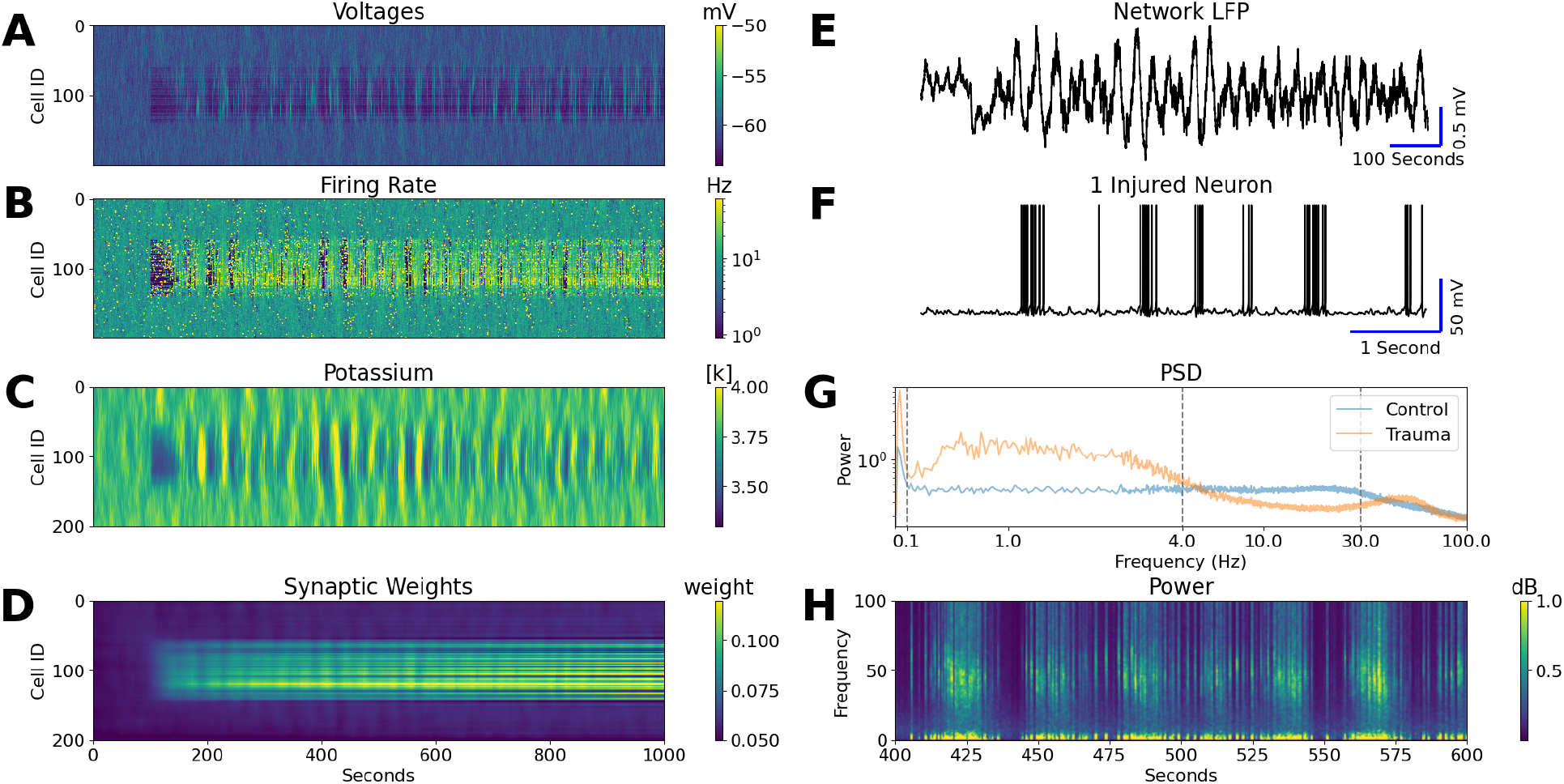
Dynamics of a network with traumatic injury inflicted at 100 seconds. Cell voltage traces (A), firing rate (B), potassium (C), and synaptic weights (D) all reveal dramatic changes that occur in the Injury Zone after the trauma is applied: voltages drop, firing rate increases, potassium shows large oscillations, and synaptic weights increase. The network LFP (E) shows much larger fluctuations than is seen in the control model, while the single neuron trace (F) now shows bursting rather than tonic firing. A comparison of PSD’s in (G) shows several key regions of change between control (blue) and trauma (orange): the trauma model has increased power < 0.1 Hz, < 4 Hz and 30 - 100 Hz, but decreased power 4 - 30 Hz. (H) shows the power spectrum over 200 seconds post injury, where the infra slow oscillations can be seen as rhythmic broadband power increases. Within and between these oscillations, short bursts of increased delta & gamma power can also be seen.

We then computed the power spectrum of neurons in the Injury Zone. Compared to the power spectrum of the same neurons in the control model, we found several important changes. The first are two distinct increases in power in the infra slow (<0.1 Hz) and Delta (0.1 Hz to 4 Hz) ranges. We then see a decrease in power from 4 Hz to 30 Hz, followed by another increase in the high Gamma range (30 Hz to 100 Hz) (Fig. 5G). A spectrogram over time shows both infra slow oscillations in broadband power, as well as shorter bursting events of high Delta and Gamma power (Fig. 5H). This mimics the bursting events seen in experimental data (Fig. 2B). While the experimental data does not show high Gamma accompanying the Delta bursts, this can be attributed to the spatial resolution of MEG; when signals from multiple cells are averaged before computing a power spectrum in our model (as is inherently done in MEG recordings), the calculated Gamma signal very quickly degrades while the Delta signal remains intact (Fig. 5-1).

These changes can be understood in the context of two prominent changes in the spatio-temporal network dynamics: increase in Infra Slow Oscillations (ISOs), and emergence of High Frequency Oscillations (HFOs). The ISOs can be seen as large fluctuations in the LFP, that is mirrored in the potassium concentration (Fig. 5E). The HFOs can be seen as very short bursts of high firing that synchronize across locally connected neurons (Fig. 6C). The locally synchronized HFOs sometimes spread through nearly the entirety of the Injury Zone, but did not significantly spread into the Healthy Zones (Fig. 6 A-B).

**Figure 6:**
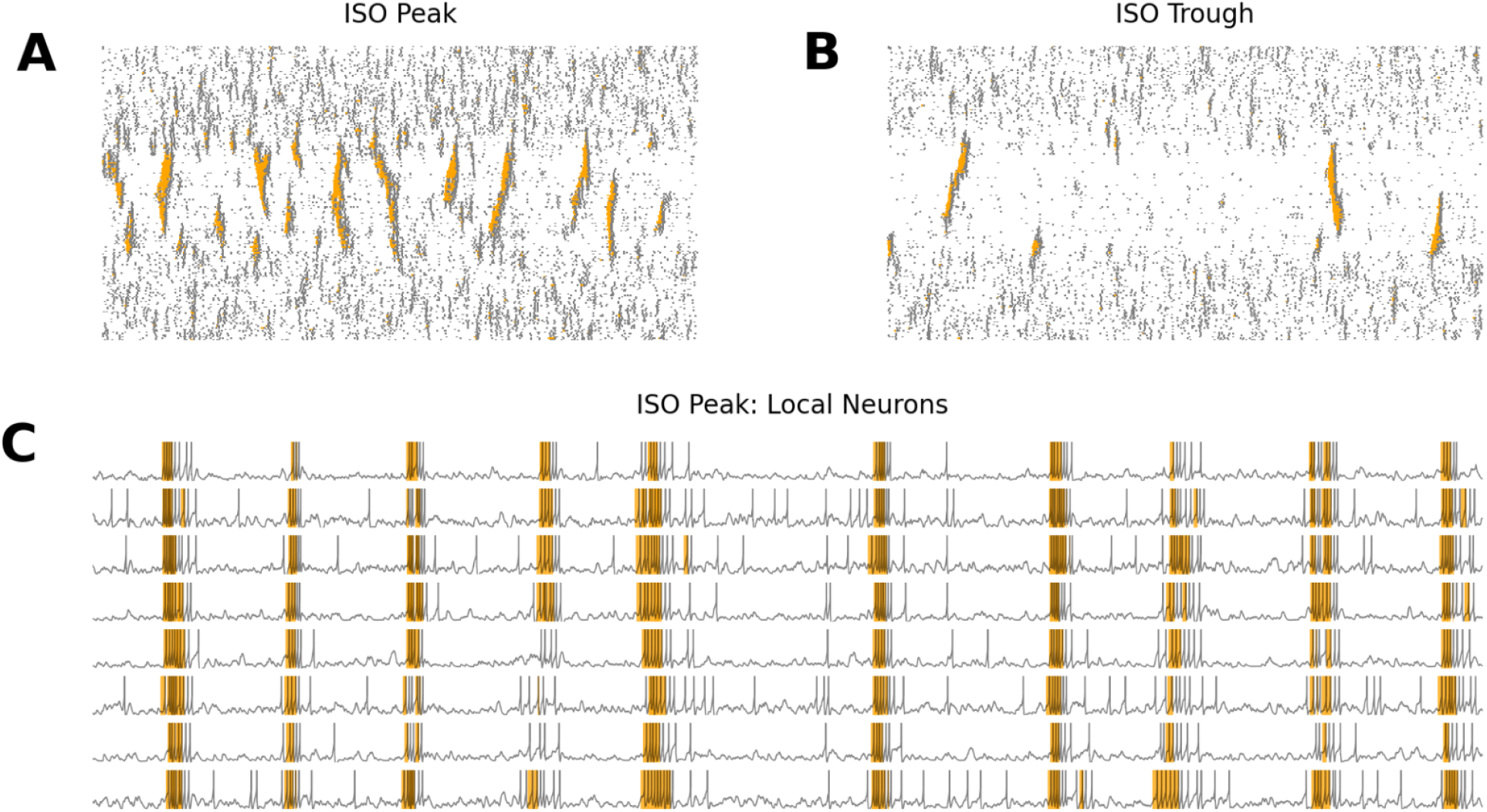
Nesting of infra-slow oscillations and HFO. Network activity during 10 seconds of a peak (A) or trough (B) of ISO, where non-HFO spikes are plotted in gray while HFO spikes are plotted in orange. HFOs were defined as 3 consecutive spikes with greater than 30 Hz frequency, and increase in density during the ISO peak. (C) Eight adjacent neurons’ voltage traces during an ISO peak, with HFO’s highlighted in orange. HFO’s occur with approximately 1 Hz frequency (10 HFO’s in 10 seconds).

ISOs and HFOs further interact with each other; the raster plots in Figure 6 taken from the peak of ISO vs the trough of a ISO show that HFOs were significantly more prominent during the ISO peaks. The HFOs are responsible for the increase in Gamma power, while the inter-HFO timing is responsible for the increase in Delta power. The loss of power in the 4-30 Hz range comes from the loss of normal tonic firing patterns.

To further investigate the origin and characterize the infra slow and high frequency oscillations, we manually manipulated the extracellular potassium levels and synaptic weights between pyramidal neurons to determine the effects of each.

### EFFECT OF POTASSIUM DYNAMICS ON NETWORK OSCILLATIONS

We began by freezing the extracellular potassium levels of the model at either a high, middle, or low point in the average potassium oscillation. Pausing at an extreme triggered HSP leading to a compensatory reaction in the synaptic weights - if potassium was paused at a high point, the weights decreased, and vice versa (Fig. 7-1). To isolate the effects of different levels of potassium, we additionally froze the synaptic weights from further updates at the same time as the potassium levels, approximately 550 seconds after the injury has occurred (Fig. 7).

**Figure 7:**
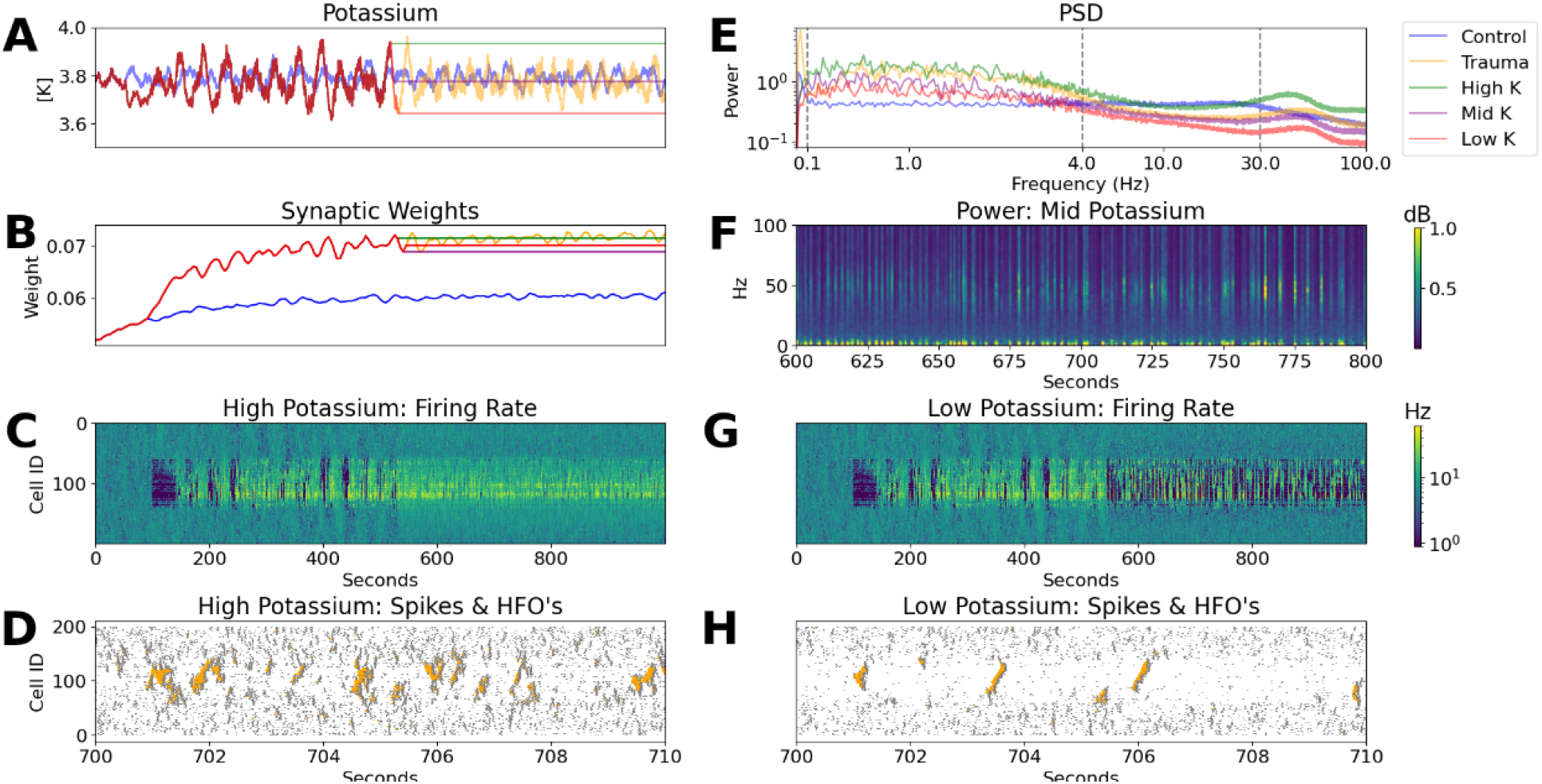
Effect of extracellular Potassium concentration on network dynamics. Potassium is frozen at a high, mid, or low point in the native fluctuation approximately 550 seconds into the injured network simulation (A). At the same time, synaptic weights are frozen (B). (E,F) A comparison of PSD’s shows a broadband power increase for high potassium (green), and a broadband power decrease for low potassium (red) compared to the original trauma case (orange). Mid potassium (purple) falls in between. (C,D and G,H) The bottom 4 plots show cell firing rates and raster plots with HFO’s identified in orange for High Potassium (C & D) and Low Potassium (G & H). High potassium shows denser HFO’s and higher firing rate, while low potassium shows the opposite.

Freezing potassium at its highest point resulted in an increased firing rate across the entire network, although the highest firing remained localized to the Injured Zone (Fig. 7C). Visualization of HFOs via a raster plot showed activity very similar to that at an ISO peak in the original injured model, with a large number of HFOs appearing densely throughout the Injured Zone only (Fig. 7D). As expected, freezing potassium at a low point resulted in the opposite: decreased firing throughout the network (although most extreme in the Injured Zone) compared to control (Fig. 7G), and a lower density of HFOs akin to what was seen in an ISO trough of the original trauma model (Fig. 7H).

We further found that freezing potassium at its lowest point resulted in a broadband loss of power; freezing at a high point in potassium resulted in a broadband increase in power; freezing at a middle level, unsurprisingly, fell in the middle (Fig. 8B-D). There is one notable exception: in all cases, there was a dramatic loss of ISO power (<0.1 Hz) compared to the base trauma case (Fig. 8A). While there was still a Fair correlation between potassium level and <0.1 Hz power, these differences were dwarfed by the decrease in power compared to the trauma network where potassium varied freely. Despite having dramatically different densities of HFOs, the peak frequency in the Gamma range (30-100 Hz) had only a Fair correlation with potassium level; notably, these shifts all occurred within the range of peak Gamma frequencies seen in the original injury model (Fig. 8E). This implies that potassium levels may narrow the distribution of intra-HFO frequencies, but do not intrinsically shift it.

**Figure 8:**
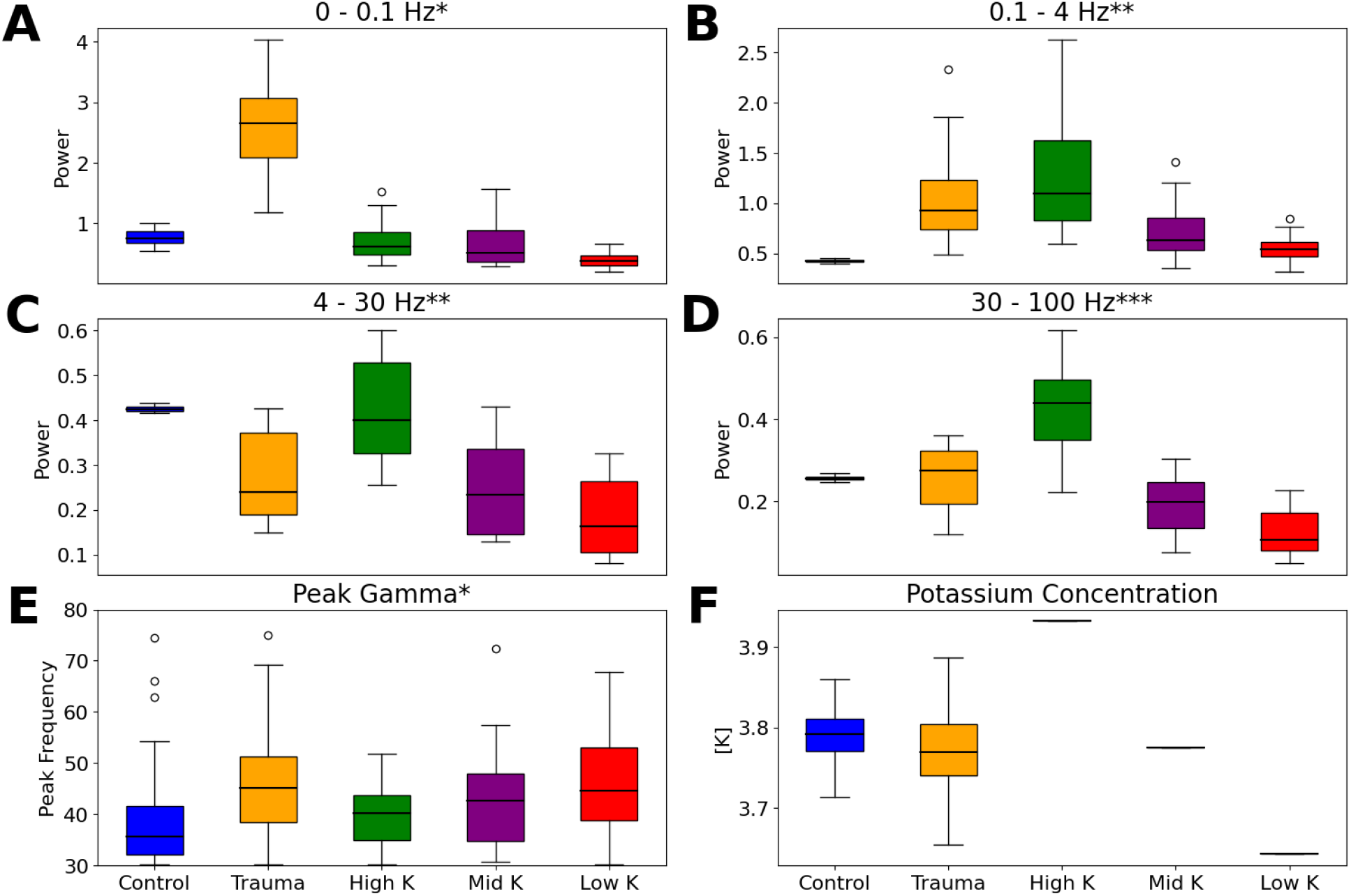
Quantification of power in several different ranges in the control (blue), trauma (orange), High K (green), Mid K (purple) and Low K (red) cases. Measurements are only taken after 600 seconds (50 seconds after potassium levels have been frozen). HSP was blocked to prevent synaptic updates. Note, there are Fair correlations between potassium level to 0-0.1 Hz power and peak Gamma power; however, these values are both retained well within the range of the control (0-0.1 Hz) or base trauma (peak Gamma) ranges. 0 - 0.1 Hz power is lost nearly uniformly as a result of the loss of potassium fluctuations. 0.1 - 100 Hz power bands all show a Moderate to Very Strong correlation, showing the broad nature of Potassium related power increases. There is no variability in potassium concentration in simulations where potassium has been frozen. */**/*** indicate a Fair / Moderate / Very Strong correlation (p<0.5) between the 3 levels of potassium.

Overall these manipulations revealed that, in agreement with our previous study of ISO in a healthy model(Krishnan et al., 2018), it is the oscillation of potassium that controls the infra slow oscillations, rather than the level of potassium. This is seen in the ablation of <0.1 Hz power after potassium is frozen at any level. We further concluded that the main effect of differing potassium levels is non-specific broadband shifts in power and in the density of HFOs (Fig. 7, D & H), rather than the characterization of the HFOs themselves (as measured by peak Gamma frequency).

Analysis of the network with exclusively High Potassium also gives an insightful characterization of the network pathology in the sense that two different states (ISO peak & ISO trough) are not being averaged together, as is the case in spectral analysis of the original trauma network. In particular, there is a much clearer increase in Gamma power over the uninjured control model (Fig. 7E, Fig. 8D) since there is no ISO trough to wash out the effects of an ISO peak.

### EFFECT OF SYNAPTIC WEIGHTS ON HFOs

To delve further into the causes of HFOs, we turned off HSP and varied the strength of synaptic weights up and down from the level natively seen after trauma (Fig. 9). This manipulation was only done on neurons in the Injury Zone. For the same reasons we froze synaptic weights at their native level in the potassium manipulations (Fig. 8), we here froze potassium at a midpoint (the same midpoint shown in Figures 7 & 8) while the weights were manipulated to isolate the effects of each. Since potassium was frozen, we again saw a dramatic loss of <0.1 Hz power that dwarfs the Fair correlation between <0.1 Hz power and frozen weight values (Fig. 9A). The ranges of 0.1-4 Hz (inter-HFO) and 30-100 Hz (intra-HFO) showed Moderate correlations (Fig. 9B,D), but there was no correlation between weight values and power in the 4-30 Hz range. This suggests that the strength of synaptic weights strongly & specifically affects the HFOs. Furthermore, there was a Moderate correlation between synaptic weights and peak Gamma frequency (Fig. 9E), indicating that weights further affect the frequency of firing within an HFO, increasing the peak Gamma range in a way that was not seen with changes in potassium levels.

**Figure 9:**
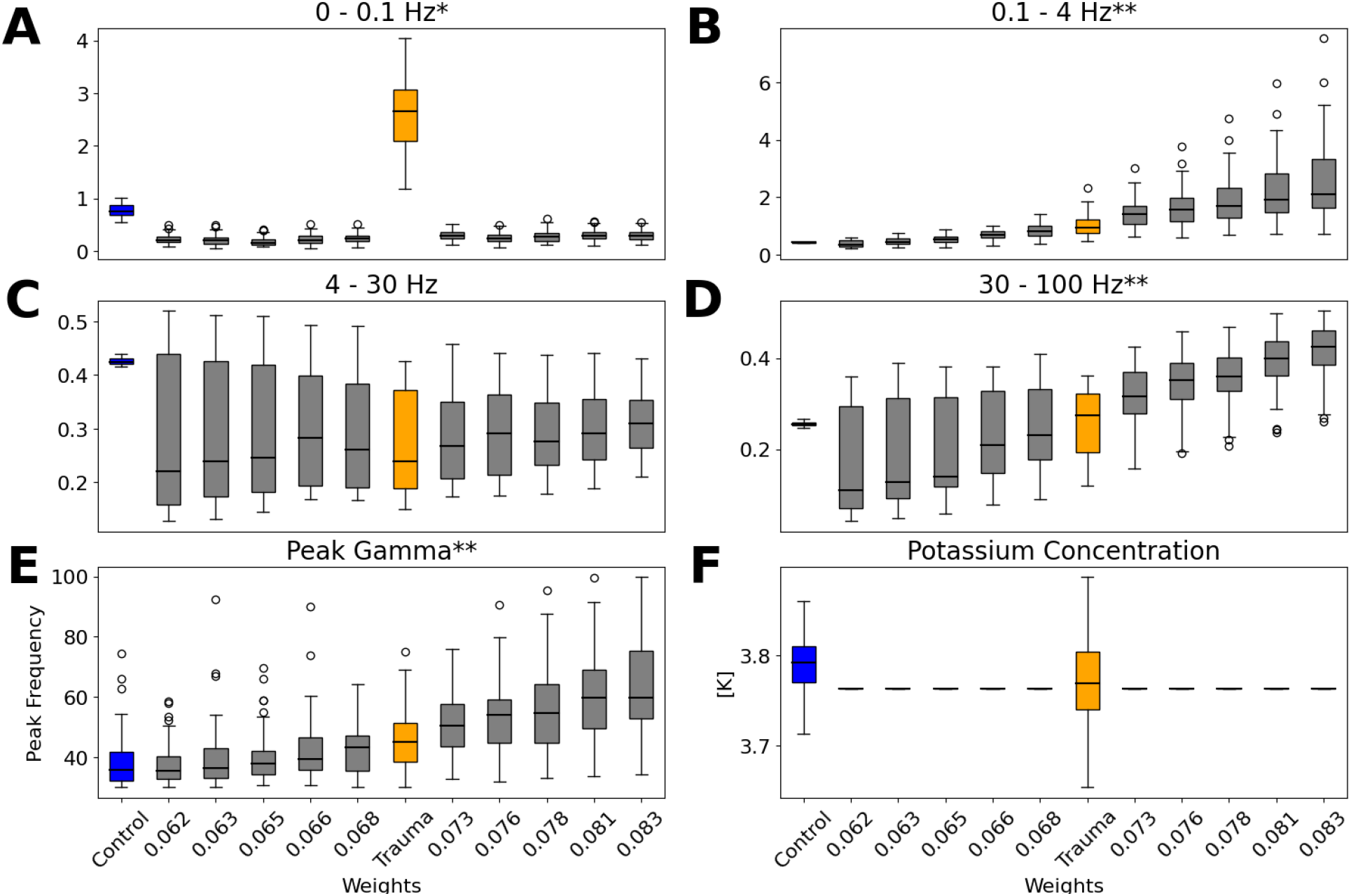
Quantification of power in several different frequency ranges as synaptic weights are manually increased or decreased after injury. Note, increase in Delta (B) and Gamma (D) power bands, as well as increase in peak Gamma frequency (E), as weights increase. (F) shows there is no variability in potassium in simulations where the weight and potassium have been manually set. */**indicate a Fair / Moderate correlation (p<0.5) of power across weight values.

In simulations where potassium was allowed to vary freely as the weights were frozen and set at different strength as in Fig. 9, we found that decreasing weights correlated with decreasing variability in the potassium concentration and increasing weights triggered seizure-like events throughout the network (Fig. 7-1). To further investigate seizure initiation and pathology, we next implemented a more severe injury that directly initiates seizures without manual manipulations of synaptic weights.

### HFOs DRIVE SEIZURE INITIATION

To induce more severe trauma, we preserved the number of injured neurons and percent of input lost (50%) to each injured neuron. However, we changed the spatial pattern of the 40 injured neurons to be 100% concentrated in the center of the network rather than interspersed with healthy neurons.

After approximately 1,300 seconds of simulation time, the first spontaneous seizure-like event was initiated in the Injury Zone (Fig. 10); two more quickly followed. The spatio-temporal pattern of seizure-like events was different between ictal events, but all are accompanied by sustained high voltages and potassium levels (Fig. 10A-B). The first seizure spreads throughout the whole network, while the second one remains largely local to the Injury Zone. After the second seizure, however, the ictal activity was initiated and spread entirely within the Healthy Zones (Fig. 10D-F). It suggests that once the ictal event has substantively spread outside of the Injury Zone, it can be maintained even in previously healthy parts of the network. The network potassium levels can be seen to increase dramatically during the seizure events (Fig. 10E), as reported in our previous computational and experimental studies(Filatov et al., 2011; Frohlich et al., 2010; Krishnan et al., 2015b, 2013; Krishnan and Bazhenov, 2011).

**Figure 10:**
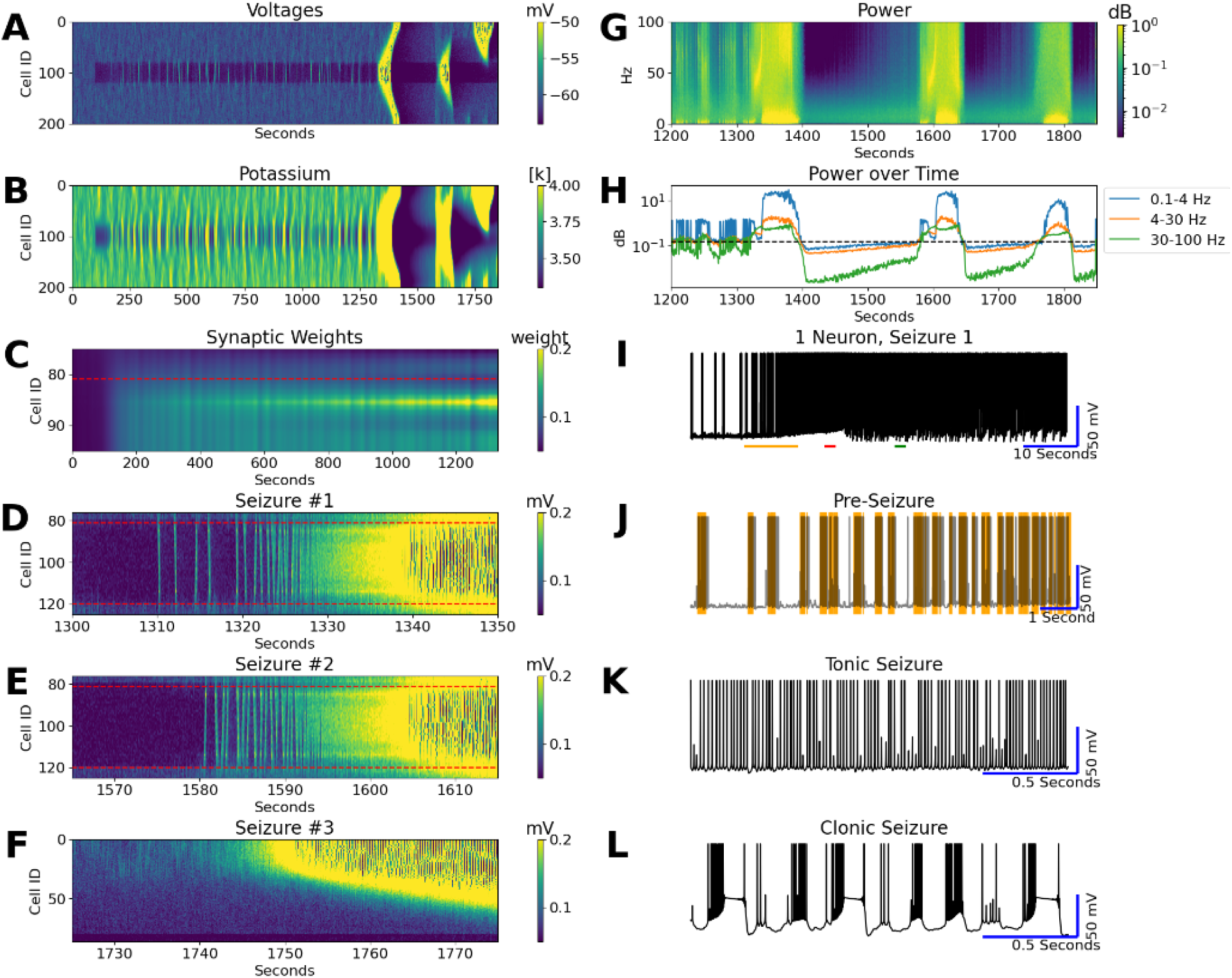
Spontaneous seizure initiation. Cell voltages (A) across the entire simulation (A). Potassium increases strongly during seizures but decreases during postictal depression (B). Synaptic weights surrounding one injury border from time 0 until just before the first seizure initiates (C). Cells just inside the injury zone (approx. cell 85) can be seen to have the highest weight increase, while cells on either side show an initial increase followed by progressive decrease. Further plots show cell voltages around the initiation time of 3 main seizures (D-F). A spectrogram of the first seizure in (G) shows a dramatic increase in ∼ 40 Hz gamma band near seizure initiation. (H) shows mean power in 3 frequency ranges (0.1-4 Hz, 4-30 Hz, and 30 - 100 Hz) over the same time period as (G). A single cell trace of the first seizure (I) shows 3 unique stages: the buildup of HFO events before seizure initiation (orange line, J), tonic seizure phase (red line, K), and clonic seizure phase (green line, L). Red dashed lines indicate injury zone borders.

Next we analyzed the initiation of ictal events. Specifically, the neurons approximately 5 neurons away from the border on either side of injury can be seen to initiate both the first and second seizure; these neurons can also be seen to have the most dramatic synaptic weight increases, while injured cells closer to the border show a weaker initial increase followed by a progressive decrease (Fig. 10C). Recall that each neuron has a connection (and local HSP calculation) radius of 5; hence, these neurons are the first within the injury region to have every cell in their connection radius also within the Injury Zone. By contrast, cells closer to the border show less dramatic HSP due to their connections with healthy cells; these already active healthy cells also respond to the injury with increases in synaptic weights, quickly compensating for lost activity in the injured cells. This resistance of the border injury cells to full recovery due to compensation of the healthy cells triggers even more extreme HSP in their immediate injured neighbors without connections to healthy cells. These cells with the most extreme increases in synaptic weights then become our seizure initiator cells.

A spectrogram of the first seizure shows a spike in Gamma, followed by substantial Delta power (Fig. 10G); this can be seen more clearly in the average power band traces (Fig. 10H). Analysis of single cell voltage traces revealed that the first ictal event was preceded by increasingly frequent HFOs (Fig. 10J). In subsequent ictal events, the network immediately transitioned to the next seizure after the postictal depression was recovered.

Single cell voltage traces (Fig. 10I-L) further revealed the typical tonic-clonic pattern(González et al., 2015; Krishnan et al., 2013) of the seizure-like event. During clonic phase neurons revealed transient depolarization blocks (Fig. 10L). This effect was observed across all neurons, injured & healthy, & regardless of where the seizure initiated. Termination of the ictal event was followed by an extended period of hyperpolarization(Krishnan et al., 2013).

## Discussion

Traumatic brain injury (TBI) is an umbrella term that covers many different patterns and causes of injury(Saatman et al., 2008), which can affect up to 1.5 million people in America alone each year(“Report to Congress,” 2019). The link between TBI and epilepsy is well known(Avramescu and Timofeev, 2008; Chauvette et al., 2016; Dinner, 1993; Timofeev et al., 2013; Volman et al., 2011b, 2011a), but the underlying causative network pathology is not well understood. However, infra slow oscillations(Franke et al., 2016; Guerriero et al., 2022; Huang et al., 2019), Delta power(Davenport et al., 2022; Safar et al., 2021; Sandsmark et al., 2017; Thomasy et al., 2016), and high frequency oscillations(Bailey et al., 2017; Culic et al., 2005; Huang et al., 2020, 2019) (HFOs) have all been shown to increase following TBI. In particular, high frequency power increases local to the seizure foci have been shown to precede ictal activity(Alarcon et al., 1995; Arroyo and Uematsu, 1992; Traub et al., 2001; Wendling et al., 2016; Worrell et al., 2004). Our biologically realistic computational model, implementing ion concentration dynamics and homeostatic synaptic scaling, natively showed all of these phenomena, and allowed us to delve deeper into their underlying causes, interactions, and roles in seizure generation.

While the base model we used in our study was a standard Hodgkin-Huxley based network of excitatory and inhibitory neurons, two unique features were critical to model post-traumatic brain dynamics – homeostatic synaptic scaling (HSP) and ion concentration dynamics. HSP maintains brain excitability at normal levels(Volman et al., 2012). Silencing a cortical culture network for two days up-regulates synaptic excitability, while an increase in activity down-regulates excitatory synaptic efficacy(Murthy et al., 2001; Turrigiano et al., 1998; Watt et al., 2000), but not all connections(Kim and Tsien, 2008). Prolonged enhanced activity induced by the blockade of synaptic inhibition or elevated [K+]o reduces the size of mEPSCs(Leslie et al., 2001; Lissin et al., 1998; Turrigiano et al., 1998). In contrast, with activity blockade, mIPSCs are scaled down, while mEPSPs are increased. After a chronic blockade of activity, intrinsic Na+ currents increase and K+ currents decrease in size, resulting in an enhanced responsiveness of pyramidal cells(Desai et al., 1999). Thus, HSP controls neuronal activity through the intrinsic and synaptic mechanisms(Murthy et al., 2001; Turrigiano et al., 1998). HSP also occurs in vivo(Desai et al., 2002).

We modeled TBI as removing afferent connections to part of the network; this reduced activity and triggered HSP that modified synaptic strength. The 2d feature of the model – ion concentration dynamics – allowed us to link activity of the network to its excitability. Reversal potentials of the intrinsic currents are usually assumed to be fixed in computer models. While an approximation even in the normal brain, this assumption becomes incorrect in the pathological brain where intense spiking may lead to significant changes of the ion concentrations and affect reversal potentials and excitability. Indeed, substantial fluctuations of extracellular K^+^ concentration ([K^+^]_o_) were found during electrically- or pharmacologically-induced paroxysmal activity(Heinemann and Lux, 1977). Extracellular Ca^2+^ concentration ([Ca^2+^]_o_) reduces during epileptic seizures, thus decreasing reliability of synaptic transmission and increasing intracellular excitability(Seigneur and Timofeev, 2011; Somjen, 2002). Extracellular Na^+^ ([Na^+^]_o_) decreases during epileptiform activity(Dietzel et al., 1982; Kraig and Nicholson, 1978; Meyer et al., 1961; Somjen, 2002), which likely corresponds to an increase of [Na^+^]_i_. Depolarizing effects of GABAa receptor activation during seizure suggest elevated levels of intracellular Cl^-^ concentration ([Cl^-^]_i_)(Cohen et al., 2002; Timofeev et al., 2002). The changes of the ion concentrations can have profound effects on the network dynamics and may be responsible for the characteristic patterns of electrical activity observed during seizures.

We used this model to investigate the underlying causes and interactions of three types of pathological network behaviors seen after TBI: (a) an increase in infra slow oscillations (ISOs), (b) the emergence of high frequency oscillations (HFOs), and (c) an increase in Delta. ISO and Delta bursts were explicitly shown in our experimental data; while increases in Gamma were less obviously seen in vivo, we have shown how sensitive high frequency oscillations can be to spatial resolution (which MEG recordings lack). We found the existence of potassium oscillations to be necessary for the existence of ISOs, but the level of potassium (low vs high) was causative only of broadband power changes. High potassium saw a greater number of HFOs and could narrow the distribution of peak Gamma (intra-HFO) values, but not notably shift it outside the range established in the original trauma network. The strength of synaptic weights, however, was tied to power changes specifically in the inter- and intra-HFO ranges; synaptic weight increases further increased the peak Gamma distribution, indicating a change in the frequency of firing within HFOs. At the highest level of synaptic weights, the mean peak frequency shifted to nearly 60 Hz with some cells reaching peak frequencies as high as 100 Hz. The HFOs varied in spatio-temporal pattern – some remained only locally synchronized while others spread through the entire Injury Zone, but rarely entered the Healthy Zone. Importantly, HFOs increased in density directly before initiation of spontaneous ictal events resembling classical spike-and-wave seizures.

In the concentrated injury model that natively generated seizures, we found, consistent with in vivo results(Chauvette et al., 2016; Timofeev et al., 2013), the internal border regions of the injury to be most prone to seizure initiation - specifically, the first cells within the injury zone to be fully connected to only other injured neurons. These cells show the most dramatic increases in synaptic weights due to HSP, and these maximally strengthened connections are most susceptible to initiating pathological behavior. This pattern of behavior can be generalized to point to the border regions of injuries as more prone to seizure initiation. After one seizure had occurred, however, we showed that the pathology can spread and seizures initiate even in previously healthy parts of the network.

The role of HSP and ion concentration dynamics in epileptogenesis and spatio-temporal patterns of epileptic seizures has been described in our previous works. Specifically, we have identified an increase in intracellular sodium and extracellular potassium levels preceding ictal events, with a marked drop in potassium during post-ictal depression(Bazhenov et al., 2004; González et al., 2018; Krishnan et al., 2015b; Krishnan and Bazhenov, 2011). The Na/K pump was shown to be crucial to maintaining stable dynamics(Krishnan et al., 2015b), while an increase in extracellular potassium in only part of a network was found to be sufficient to generate oscillations throughout the network(Bazhenov et al., 2004). Further studies of the role of HSP have shown that sufficiently upregulated HSP can generate periodic slow oscillations as well as bursting activity in individual neurons(Fröhlich et al., 2008; González et al., 2019; Krishnan et al., 2013). We have then shown that age-related changes in the mechanism of HSP can explain differences in seizure susceptibility in older populations(González et al., 2015; Timofeev et al., 2013), and that spatially compact trauma is more prone to epileptic activity than diffuse trauma due to differential HSP mechanisms(Volman et al., 2011b, 2011a). The novelty of our new study here is linking these two mechanisms of HSP and ion concentration dynamics to an increase in delta power and HFO events after TBI.

Our model does not stand in isolation. Fink demonstrated the existence of both normal and abnormal HFOs, produced by different mechanisms(Fink et al., 2015). Importantly, they further showed that HFOs can be generated by a variety of different network structures, just as we see the same types of HFOs appearing in our two different models of TBI. This lends support to the idea of generalizable effects of TBI despite potentially dramatically different causes and patterns of injury. Molaee-Ardekani further identifies”resonance” modes as necessary to generate high frequency outputs; these modes are at the border of stable (normal firing) and unstable (ictal firing) regions(Molaee-Ardekani et al., 2010). This lends further understanding to the role the HFOs may play in transitioning between healthy & pathological behavior. Finally, several models put forward by Traub explore the cellular mechanisms of HFOs, specifically pointing to the coupling of pyramidal neurons via gap junctions(Traub et al., 2010, 2002; Traub and Bibbig, 2000). As a whole, our own conclusions in this paper both support and are supported by these results.

The effects seen natively occurring in our model following injury closely resemble effects seen in our (and many others) experimental data. Given the complex ion dynamics of our neurons that are modeled directly after in vivo processes, it stands to reason that our simulations are as reliable and relevant to true biological physiology as is currently possible. Our study is another step towards a deeper understanding of the effects of TBI, and how they may lead to epileptic activity. This model further creates a testing ground for potential interventions following TBI that address the underlying pathology in the long term.

## Acknowledgments

Supported by NIH (grant R01NS104368 to MB)

