## Supplemental Figures for "Network effects of traumatic brain injury: from infra slow to high frequency oscillations"

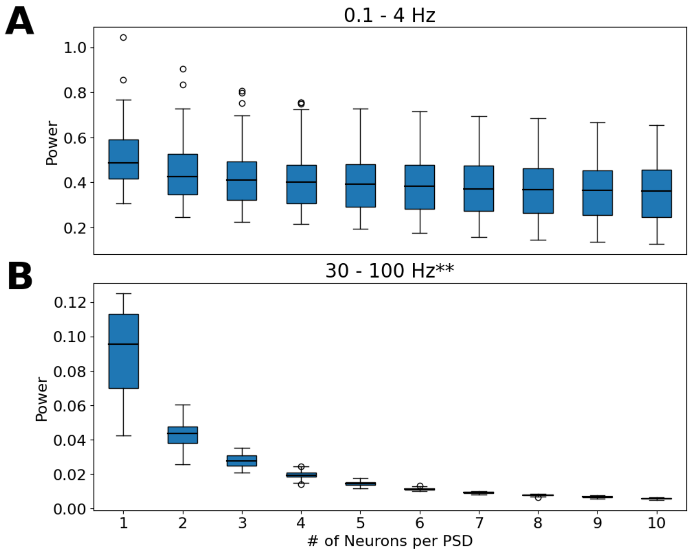


Figure 5-1: As the number of neurons averaged together before computing a PSD increases, the Delta signal (A) remains largely consistent while the Gamma signal (B) very quickly degrades and is lost.

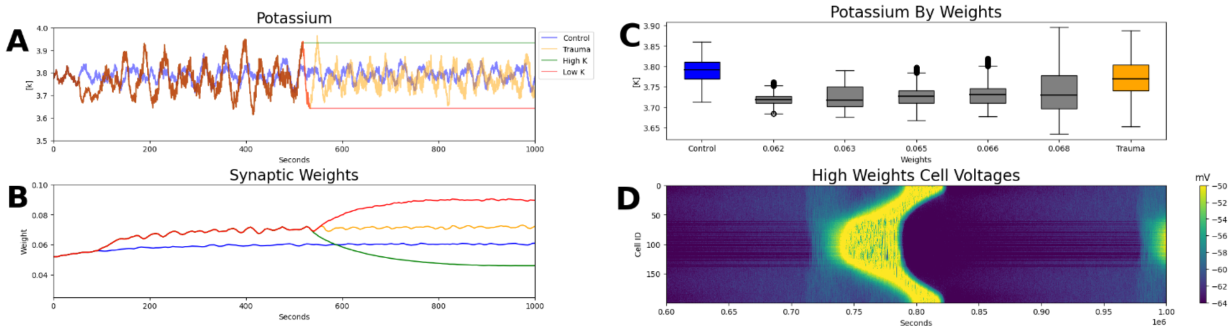


Figure 7-1: (A & B) show that when potassium is frozen at an extreme value, there is a strong compensatory reaction in synaptic weights, (C) Potassium levels and variability by synaptic weight value, (D) Network voltages when synaptic weights are increased by 10%.
